# EEG biomarkers of free recall

**DOI:** 10.1101/2021.04.30.442192

**Authors:** B. S. Katerman, Y. Li, J. K. Pazdera, C. Keane, M. J. Kahana

## Abstract

Brain activity in the moments leading up to spontaneous verbal recall provide a window into the cognitive processes underlying memory retrieval. But these same recordings also subsume neural signals unrelated to mnemonic retrieval, such as response-related motor activity. Here we examined spectral EEG biomarkers of memory retrieval under an extreme manipulation of mnemonic demands: subjects either recalled items after a few seconds or after several days. This manipulation helped to isolate EEG components specifically related to long-term memory retrieval. In the moments immediately preceding recall we observed increased theta (4-8 Hz) power (+T), decreased alpha (8-20 Hz) power (-A), and increased gamma (40-128 Hz) power (+G), with this spectral pattern (+T-A+G) distinguishing the long-delay and immediate recall conditions. As subjects vocalized the same set of studied words in both conditions, we interpret the spectral +T-A+G as a biomarker of episodic memory retrieval.

## Introduction

Our ability to recall past experiences is the hallmark of episodic memory. In laboratory studies, lists of discrete nameable items, such as words, serve as sets of mini-experiences, and the act of recall is the motor output (vocalization or typing) of the remembered items. The magic of memory retrieval happens during the moments leading up to the motor output, but this period also includes other non-mnemonic signals, such as the planning of a motor response and the activation of semantic and perceptual representations associated with the remembered item.

Prior research using intracranial EEG recordings has uncovered a network of brain regions in which increased high frequency activity and concomitant decreases in lower frequency activity mark the moments leading up to spontaneous verbal recall as compared with matched periods of silence during the recall period (Burke, Sharan, et al., 2014; Greenberg, Burke, Haque, Kahana, & Zaghloul, 2015; Solomon et al., 2017). In some cortical sub-regions and for some retrieval contrasts, low-frequency (3-8 Hz) theta activity increases during the putative retrieval period (see Herweg et al., 2016, for a review). Researchers would ideally like to use these recall biomarkers to probe the dynamics of memory retrieval in the absence of motor output. Such signals could, for example, be used to study the hypothesized role of covert retrieval in memory consolidation and other learning processes. However, the interpretation of these recall biomarkers is problematized by the concomitant presence of non-mnemonic neural processes that accompany recall even under minimal memory demands.

Comparisons of correct and incorrect recalls (intrusions) offers a potential solution (Long et al., 2017) These studies have shown, for example, that hippocampal high-frequency activity increases moreso prior to correct recalls than prior to intrusion errors. As incorrect recalls presumably involve identical, or at least very similar, motor planning activity, differences between these conditions more likely reflect mnemonic retrieval. However, intrusions may show similar signals to those of successful recall as the subject is actively engaged in recall of a word they believe to have been previously presented. As such, these contrasts may mask important neural correlates of memory retrieval that appear similarly for both true and false memories.

The present study sought to elucidate the biomarkers of episodic memory retrieval by examining recall of words under an extreme manipulation of retrieval demands: In an immediate recall condition, subjects recalled a single just-presented word after a brief delay. In a long-delayed recall test, subjects attempted to recall items learned across multiple days prior to the test day (i.e., a minimum of 16 hours prior to the recall period). Over the course of ten experimental sessions administered on different days, subjects contributed data in each of these conditions.

## Method

### Subjects

Fifty-seven young adults (ages 18-35, 24 reported male, 29 reported female, 53 reported right-hand dominant), recruited among the students and staff at the University of Pennsylvania and neighboring institutions, each contributed 10 sessions of immediate word recall data and five sessions of delayed recall data. All subjects provided informed consent to participate in our study, which was approved by the Institutional-Review Board at the University of Pennsylvania. Subjects were excluded (*n*=17) if they did not complete at least 7 sessions or if they had an average rate of recall lower than 70% during the immediate recall task. The data reported here came from Experiment 5 of the Penn Electrophysiology of Encoding and Retrieval Study (PEERS). This is the first paper to report data from PEERS-Experiment 5.

### Data Availability

All PEERS data, including the full dataset reported and analyzed in the present manuscript, may be freely downloaded from our public repository http://memory.psych.upenn.edu/data. Analysis code for this manuscript is also available at the same URL.

### Experimental Task and Behavioral Analyses

Each of the first five sessions (henceforth, *Phase I*) consisted of one block of 10 practice trials, followed by 24 blocks of 24 trials each. Each block began with a 10-second countdown. After the countdown was complete, the first trial of the block began. On each trial, a black screen was shown for a jittered 1000--1600 ms (uniformly distributed), after which a single word appeared onscreen in white text for 1200--1800 ms (uniformly distributed). Following presentation, the screen went blank again and subjects were instructed to pause briefly, and then vocalize the word they had just seen. If they began speaking within 1.0 seconds of word offset, the message “Too fast.” appeared on the screen in red text. By avoiding these messages subjects could increase the size of their bonus payment. After the subject finished speaking, a tone sounded, marking the end of the current trial. Speech was detected using a volume amplitude threshold. In addition to the 10-second countdown between blocks, two 2-minute mid-session breaks were administered after block eight and block 16.

Phase II of the experiment began on the day of the sixth session and continued to the final session of the experiment. In Phase II the practice block and 24 experimental blocks were preceded by a 10-minute initial externalized free recall period. Subjects were instructed to recall as many words as possible from the previous sessions in any order, while also vocalizing any additional words that come to mind in their attempt to recall these items (e.g., Kahana, Dolan, Sauder, & Wingfield, 2005; Lohnas, Polyn, & Kahana, 2015; Zaromb et al., 2006). Our lists comprised the same 576-word pool as used in the PEERS4 study (Aka, Phan, & Kahana, 2020; Kahana, Aggarwal, & Phan, 2018; Weidemann & Kahana, 2020). Subjects saw the same 576 words in each of their 10 sessions, but the ordering of these words was randomized for each session. Although subjects saw the 576 words across multiple sessions, the only information identifying these words as belonging to the target list was their occurrence within the context of our experiment, thus making this a test of long-term episodic memory.

### EEG Post-Processing and Spectral Decomposition

We recorded electroencephalographic (EEG) data using a 128-channel Biosemi system with a 2048 Hz sampling rate. We applied the following preprocessing steps to the data from each session. First, we applied a 0.5 Hz highpass filter to eliminate baseline drift. We then partitioned the recording into thirds by splitting at the end of each mid-session break. For each partition of the data, we identified bad channels as those with extremely high or low (|*z*|>3) log-variance, or which had an extremely high (*z*>3) Hurst exponent relative to other channels. We identified bad channels separately for each partition, as problematic electrodes were often corrected during mid-session breaks.^1^ We then dropped the bad channels from their respective partitions and applied a common average reference scheme. We next performed independent component analysis (ICA) on each of the three partitions to decompose the data into 128-(*n*+1) components, where *n* is the number of bad channels that were dropped, and used the localized component filtering method of DelPozo-Banos and Weidemann (2017) to filter artifactual time points from the components. Data points were identified as artifacts if they exceeded that component’s interquartile range by three times the magnitude of that range. We then reconstructed the original channels from the cleaned components, interpolated bad channels using spherical splines, and applied a fourth-order Butterworth notch filter at 60 Hz to eliminate electrical line noise.

We used a multitaper method (MNE-Python software package; Gramfort et al., 2013, 2014) to estimate spectral power at each electrode over 4--128 Hz and log transformed the resulting signal. Electrodes were grouped into regions of interest (see Figure 2) and the corresponding powers averaged for each frequency. We spaced frequencies every 2 Hz in the range of 4--26 Hz, and every 6 Hz within 26--128 Hz, resulting in 29 frequencies of interest. We avoided the Morlet wavelet method, as convolving low frequency wavelets using buffer periods may allow speech artifacts to intrude in the power estimates of intervals just prior to subjects’ vocal responses. We used a 500 ms moving window centered at multiple time-points relative to the start of recall events (i.e., vocalization of a recalled item), with a 50 ms step size. To minimize potential artifacts from pre-motor activity, we extracted spectral patterns in a 500 ms window ending 250 ms prior to the annotated time of speech onset unless otherwise noted. We excluded from our analysis recall events that occurred within 1500 ms of the onset of the prior recalled item. In addition to the interval preceding vocalization, we identified 1000 ms “deliberation” periods of silence during the delayed recall test that did not overlap with a preceding vocalization (i.e., within 500 ms of vocalization onset) or a subsequent retrieval interval of interest. Deliberation periods for both immediate and delayed recall contrasts were matched to recall events using linear sum optimization, which minimizes the total time difference between recall events and matched deliberation periods, and were constrained to events between the first and last delayed recalls within a session.

**Figure 1.**
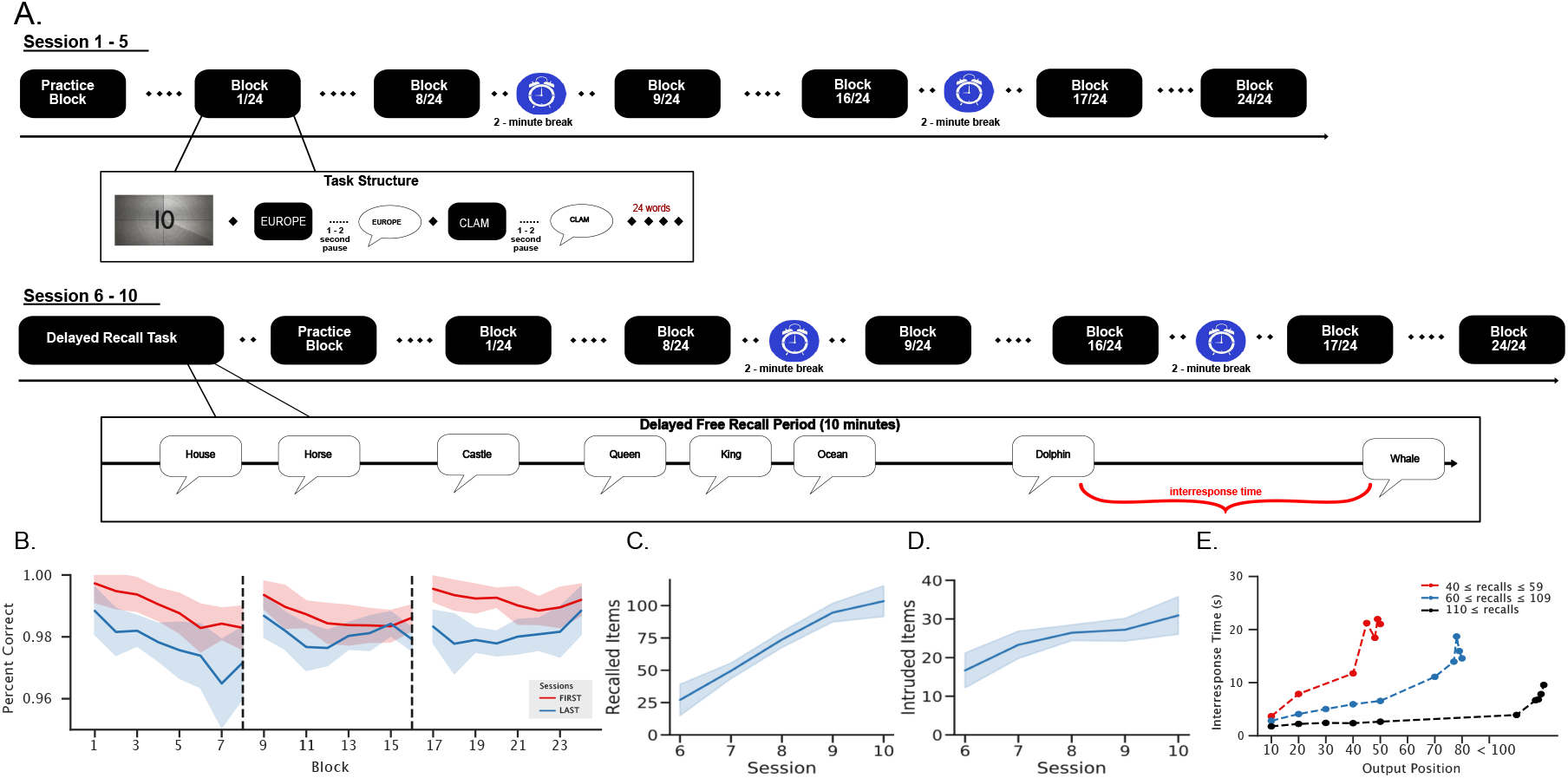
Experimental Paradigm and Behavioral Data. **A.** During Sessions 1-5, subjects performed an immediate recall task for each of 576 words, silently reading a word which they verbally recalled after waiting approximately 1 second. On average, subjects responded 1.53 s after word onset. Each of sessions 6-10 began with subjects attempting to freely recall the 576 words that they had seen on each of the preceding sessions along with any other words that come to mind during a 10 minute retrieval interval. Subjects then performed the same immediate recall task as on earlier sessions. **B.** Subjects exhibited very high levels of immediate recall across all sessions, with performance dropping modestly across blocks and recovering following breaks. **C.** Subjects exhibited modest levels of delayed recall on the first (surprise) recall test given on Session 6, but performance rose sharply across subsequent sessions, hitting an average of 103 correct recalls by the final session. **D.** Intrusions similarly rose across sessions, but much less quickly than successful recalls. **E.** Inter-response times (IRTs) increased with output position; on sessions where subjects recalled a larger proportion of items, IRTs were generally faster throughout the retrieval period but rose sharply during the last few correct recalls.

**Figure 2.**
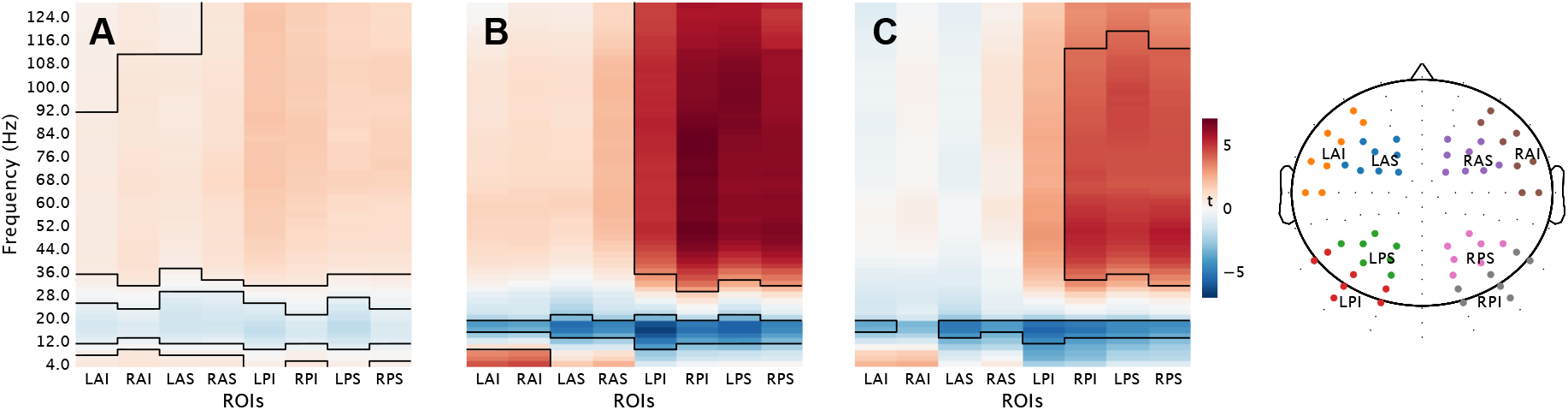
Statistical maps illustrating relative increases (red) and decreases (blue) in spectral power across key memory contrasts for eight regions of interest. **A.** Delayed Recall vs. Deliberation. **B.** Delayed Recall vs. Immediate Recall. **C.** Deliberation vs. Immediate Recall. Panels **A** and **B** use matched time periods from the first five sessions of Immediate Recall and the five sessions of Delayed Recall following those first five sessions. Black-bordered regions indicated significant FDR-corrected t-tests on within subject difference scores (*p* < .05). Color bar corresponds to t-stat differences in panels A, B, and C. Electrodes locations corresponding to each region of interest appear on a schematic view of an electrode net (LAI,RAI: left/right anterior inferior; LSA,RAS: left/right anterior superior; LPI,RPI: left/right posterior inferior; LPS,RPS: left/right posterior superior).

During delayed recall, we defined successful memory events as the memory search intervals immediately preceding recalls of list items (correct recalls). Unsuccessful memory search periods preceded recall errors (i.e., intrusions from outside of the word pool: extra-list intrusions). To identify spectral features specific to successful recall on the delayed recall test, we performed contrasts against successful immediate recall and successful delayed recall across eight regions of interest (ROI). ROIs were selected based on previous studies using similar EEG caps (Weidemann, Mollison, & Kahana, 2009). Only immediate recall events from the first ten minutes of the first five sessions were included in these contrasts; this time period is matched to the delayed recall that occurs in later sessions and only covers immediate recall events where the subject is unaware of future delayed recall tests. We also performed a contrast between immediate recall events from the first five sessions and deliberation periods from the long-delayed recall test using the same matching procedure. We aggregated desired events from each session within subjects and performed a *t*-test to produce a variance-normalized difference score for every frequency-by-ROI pair within subjects. We subsequently used two-tailed one sample *t*-tests (with FDR correction) to assess group-level differences.

## Results

Figure 1A illustrates the basic structure of the experiment. In each of 10 experimental sessions, subjects performed a simple immediate-recall task. As each of 576 words appeared individually, subjects read each word silently, and then, after a one second delay, recited the word aloud. At the start of the 6th session (phase II), subjects were given a surprise long-delay recall task: We asked them to recall as many of the 576 words as they could remember in any order as well as any words that come to mind that may have not been presented in previous sessions. We then had subjects perform the same immediate recall task as on previous sessions. Sessions 7-10 replicated the methods of Session 6 except that the initial recall test could no longer be deemed a surprise.

As expected, subjects recalled items with very high accuracy (above 96%) in the immediate recall condition. Figure 1B illustrates accuracy of immediate recall across the 24 test blocks separately for phases one and two. Accuracy fell slightly across blocks but recovered after each of the two-minute breaks (see Figure 1A), possibly due to build-up and release from proactive interference (e.g., Lohnas et al., 2015; Underwood, 1957). Figure 1C illustrates accuracy of long-delay free recall. On the first session of phase II, subjects recalled an average of 29 words (5% of the total pool of 576 words). Their performance on subsequent sessions increased dramatically, reaching an average recall rate of 103 items by the 10th session. This rate of increase (about 25 items per session) is much higher than the implied rate of learning across sessions 1-5, but this likely reflects the difference between incidental and intentional encoding.

Figure 1D illustrates the number of extra-list intrusions committed by subjects across the five sessions of phase II. Here we see that subjects committed a much larger percentage of intrusions than seen in standard within-session immediate and delayed recall tasks (Zaromb et al., 2006). However, the overall fraction of intrusions decreased over sessions, dropping from 37% on Session 6 to 23% on Session 10. Given the unusually-long delay in our free recall condition, and the very high rates of recall achieved by the 10th session, we asked whether inter-response times (IRTs) across the recall period exhibited the same pattern of growth as documented in more standard free-recall paradigms (Rohrer & Wixted, 1994; Murdock & Okada, 1970). Figure 1E shows average IRTs based on the total number of recalled words during delayed-recall sessions. This analysis demonstrates when subjects recall many items, they exhibit much faster IRT overall. However, subjects still display a sharp increase in their IRTs as they approach their final recalled item.

Our primary question concerned how neural activity, as measured through spectral analysis of EEG recordings, signaled the process of spontaneous retrieval of previously experienced items. We first addressed this question by comparing the 500 ms pre-vocalization period in the long-delay recall condition with matched periods of silence separated by at least 0.5 sec from prior and subsequent recalls (we refer to these as deliberation periods; see *Methods*). This comparison of retrieval and deliberation periods revealed increased high-frequency activity and decreased alpha-band power across most regions of interest (See Figure 2A; black outlined regions indicate statistically-significant frequency-region pairs, FDR-corrected *p*<0.05), extending previous intracranial-recording studies that identified similar retrieval biomarkers in both cued recall and free recall tasks (Burke, Ramayya, & Kahana, 2015; Burke, Sharan, et al., 2014; Greenberg et al., 2015). Whereas those earlier studies examined recall that took place within minutes of item encoding, the present study asked subjects to recall items that had not been seen (experimentally) for at least 16 hours.

At anterior electrodes we observed increased theta-band activity during the pre-vocalization memory-retrieval period. Although previous studies have frequently reported theta increases during successful recognition memory (Addante, Watrous, Yonelinas, Ekstrom, & Ranganath, 2011; Guderian & Düzel, 2005; Herweg et al., 2016; Osipova et al., 2006) most (standard) free recall studies find retrieval-related *decreases* in theta and alpha-band power (Burke, Sharan, et al., 2014; Kragel, Koban, Barrett, & Wager, 2018; Solomon et al., 2017; Solomon, Stein, et al., 2019). However, several free and cued recall studies have found mixed results, with positive effects in specific brain regions, such as right anterior temporal pole (Burke, Long, et al., 2014) and for specific contrasts, such as semantically-clustered vs. non-clustered recall transitions (Solomon, Lega, Sperling, & Kahana, 2019). Our study demonstrates retrieval-related increases in low theta power coupled with alpha-band decreases in a long-delay recall task, conditions likely to place stronger demands on associative retrieval processes (Herweg, Solomon, & Kahana, 2020).

Comparisons of pre-vocalization and deliberation intervals do not, however, uniquely identify the process of memory retrieval. This is because prevocalization periods also differ from deliberation in the presence of premotor activity related to the vocalization of recalled items. Switching to another recall modality, e.g., typing, would simply replace this confound with a different one. We therefore employed an immediate recall task as a control for motor activity that may confound our comparison between pre-vocalization and deliberation intervals.

Using the pre-vocalization period in our immediate-recall task as a control, we found that increased theta, decreased alpha, and increased HFA mark spontaneous (free) recall following a long delay (Figure 2B; black outlines indicate statistically-significant frequency-region pairs, that met an FDR-corrected *p*<0.05 threshold). This comparison recapitulates the spectral pattern observed in our comparison between long-delay recall and deliberation, but without the confound of vocalization present in Figure 2A and earlier studies. One difference, however, is that in our tighter contrast between immediate recall and long-delay recall, the retrieval related theta effect appeared to be restricted to anterior-inferior ROIs. Given the far greater demands on episodic memory retrieval in the long-delay condition, but the matched vocalization in both conditions, we interpret these biomarkers as reflecting neural correlates of context-dependent memory retrieval. Comparisons between immediate recall and deliberation (Figure 2C) further support this interpretation. Here we see that trying to retrieve after a long delay (in the deliberation periods) exhibits higher HFA and low alpha-band power than during the period immediately preceding a motor response in the immediate recall condition (black outlines indicate statistically-significant frequency-region pairs that met an FDR corrected *p*<0.05 threshold).

Because high-frequency signals provide excellent temporal resolution, we sought to examine the timing of the increased retrieval-related HFA seen prominently in posterior ROIs. Figure 3 shows the time course of high-frequency activity (HFA) leading up to retrieval in the long delay and immediate recall conditions. We illustrate this time course separately for immediate recalls contributed during phases one and two. Error bands indicate the 95% confidence interval based upon between-subject variability in high-frequency activity. For all three conditions, HFA rose in the moments leading up to recall, but the prevocalization HFA was highest for delayed recall, lower for phase-two immediate recall, and lowest for phase-one immediate recall. For comparison, we indicate the baseline gamma power in the long delay deliberation periods. This ordering of conditions aligns with the hypothesized episodic memory retrieval demands across these conditions (such demands being highest in delayed recall, and lowest in immediate recall when a memory test is not expected). Given that the expectation of a subsequent test would lead to better memory encoding during phase II, as shown in Figure 1C, mnemonic processes likely exerted a greater influence on immediate recall in phase II than in phase one.

**Figure 3.**
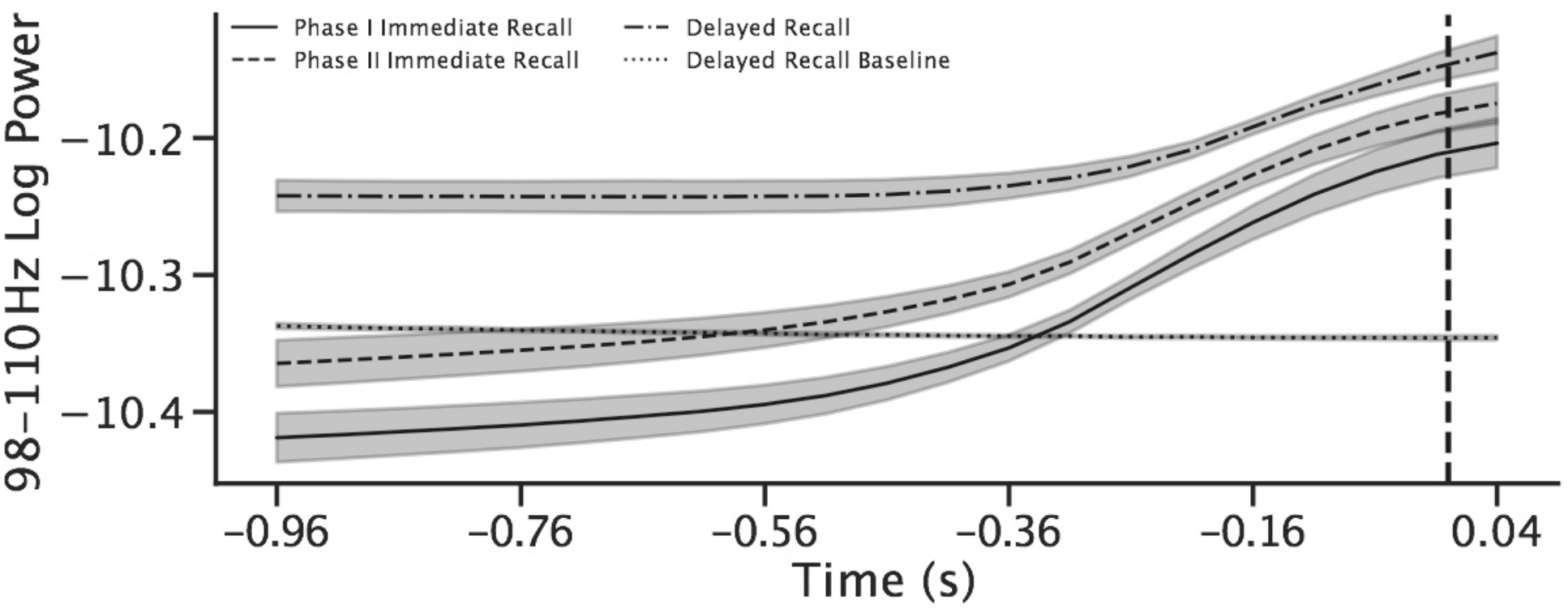
Time course of high-frequency activity leading up to correct recalls. Delayed recall, immediate recall, and delayed recall baseline log high-frequency activity (98-110 Hz power) at posterior electrodes in the 1.0 second leading up to vocalization of recalled items, averaged for all correctly recalled items and across subjects. Results are shown separately for phases one and two. Error bands reflect 95% confidence computed by the method of Loftus and Masson (1994).

## Discussion

Humans possess a remarkable ability to search their memory for previously experienced items learned in a given context. When asked to recall without the aid of specific cues, subjects generate their own retrieval cues, based upon the context at the time of test as well as the contextual representations evoked by recently remembered items (Kahana, 2020). Here we examined the electrophysiological (EEG) correlates of spontaneous memory retrieval under conditions designed to vary subjects’ reliance on contextual retrieval between two extremes: In a long-delay recall condition, we asked subjects to freely recall items not seen in at least 16 hours but encoded on previous sessions; in an immediate recall condition, we asked subjects to read a word, pause for >1 second, and then speak the word aloud. We chose these extreme contrasts to help distinguish pre-motor activity related to vocalization from context-dependent memory-retrieval processes required in the long-delay task.

Contrasting immediate recall of a single word with long-delayed free recall of the entire 576-word pool identified a spectral signature of spontaneous memory retrieval. Increased theta- and gamma-band power (4-8 Hz and >40 Hz respectively), coupled with reduced alpha- and beta-band power, marked periods immediately preceding recall of previously studied items. The 576 words that subjects attempted to recall shared one essential property: namely, that they were experienced in the context of our laboratory experiment. As such, the spectral signature of increased theta, decreased alpha and increased gamma appears to identify a neural signature of context-dependent memory retrieval.

Although our contrast of immediate recall and long-delay recall should reduce the influence of premotor and electromyographic signals in comparisons between pre-vocalization and baseline periods, we recognize that premotor activity may differ between our short- and long-delay recall tasks. Whereas the long inter-response times in our delayed recall condition (see Figure 1e) would necessitate subjects initializing motor commands prior to vocalization, subjects in our immediate recall task could have more easily anticipated and maintained these motor programs. Although other differences between the immediate- and long-delay recall conditions could also give rise to differences in neural activity, matching for word vocalization across these conditions should provide a cleaner index of memory-related neural activity.

Most prior EEG investigations of human memory have either compared encoding of subsequently remembered and forgotten items (Staudigl & Hanslmayr, 2013; Klimesch, Doppelmayr, Russegger, & Pachinger, 1996; Burke et al., 2015; Solomon et al., 2017; Hanslmayr, Spitzer, & Bäuml, 2008) or have compared successful and unsuccessful discrimination between targets and lures in a recognition task (Jacobs, Hwang, Curran, & Kahana, 2006; van Vugt, Sekuler, Wilson, & Kahana, 2013; Guderian & Düzel, 2005; Hsieh & Ranganath, 2014; Osipova et al., 2006). These, and other prior studies reviewed by Herweg et al. (2020) and Hanslmayr and Staudigl (2014), report both positive and negative correlations between theta-band power and successful mnemonic processes. In the present study we focused on the EEG correlates of free recall, a process rarely studied using non-invasive recording methods due to the presence of large electromyographic (EMG) artifacts caused by vocalization during recall. By contrasting long-delayed recall with immediate recall, we find that increased theta and decreased alpha power mark periods of episodic memory retrieval. Pastötter and Bäuml (2014) report a similar theta-alpha pattern in a cued recall task.

Our finding that memory-related theta increases appeared at the lower bound of the theta range aligns with several earlier reports using intracranial recordings to examine memory encoding (Miller et al., 2018; Lega, Jacobs, & Kahana, 2012; Lin et al., 2017) as well as scalp EEG and MEG studies of successful recognition memory (Herweg et al., 2016; Gruber, Tsivilis, Giabbiconi, & Müller, 2008). Studies reporting negative correlations between theta and successful memory encoding or retrieval either did not use contrasts that selected for the associative or contextual aspects of memory retrieval, or they averaged across a broader frequency range that would include both low-frequency increases and alpha-band decreases (see Herweg et al., 2020, for a review).

Our finding that decreased alpha band-power accompanies memory retrieval dovetails with several recent studies. Griffiths (2019) observed decreased alpha power (and increased gamma power) in two variants of an associative recognition task; one involving word and multi-modality stimuli and one involving animal image cues paired with face/place images. Griffiths (2021) additionally found decreases in alpha power during successful memory retrieval and encoding.

Simultaneous recordings of local field potentials and single neuron activity have implicated high-frequency activity (HFA) as a correlate of neuronal firing rates (Manning, Jacobs, Fried, & Kahana, 2009). Studying neurosurgical patients with indwelling electrodes, Burke et al (2015), Solomon et al (2017), and Long et al. (2018) have all found greater HFA immediately preceding recall than during matched deliberation intervals. Although invasive recordings minimize the influence of premotor artifacts, it is still likely that some non-mnemonic factors contributed to those reported results. Our finding that high-frequency activity increases in the moments leading up to spontaneous free recall when comparing long-delay with immediate recall (Figure 2B) directly implicates HFA increases in the cognitive processes involved in episodic memory retrieval.

The spectral methods employed in our study cannot distinguish between narrow-band (periodic) and broadband (aperiodic) components of the EEG signal. To determine the relative contributions of periodic and aperiodic components, Donoghue et al (2020) introduced “fitting of oscillations and one-over *f* noise,” or FOOOF method. This method estimates the aperiodic (1/*f*) component of the EEG signal, which is then subtracted from the original signal to estimate the periodic components. Wen and Liu (2016) propose a different approach to decomposing the EEG signal. In their irregular-resampling auto-spectral analysis (IRASA) method, they use mathematical properties of the aperiodic (1/*f*), or fractal, component of the EEG signal to separate these from the narrowband components. Using methods such as IRASA or FOOOF to determine the relative contributions of periodic and aperiodic signals will likely offer additional insights into the neural basis of spontaneous memory retrieval.

The present study identifies a pattern of electrophysiological biomarkers of episodic recall: increases in frontal slow-theta (+T), and decreases in alpha-band power (-A) and increased high-frequency, or gamma, activity with a posterior distribution (+G). These biomarkers, tag (+T-A+G) the process of retrieval from episodic memory. They also add to an emerging body of evidence demonstrating the utility of non-invasive methods for decoding cognitive states (Chakravarty, Chen, & Caplan, 2020; Noh, Liao, Mollison, Curran, & de Sa, 2018; Weidemann & Kahana, 2019, 2020). Given the ease of collecting non-invasive EEG data from human research subjects, and human’s potentially unique ability to spontaneously recall verbal items, the +T-A+G biomarker of episodic recall can serve as a basis for future studies that investigate the role of retrieval cues, such as the temporal, semantic and spatial contexts surrounding experienced items.

## Footnotes

1 Recordings during break periods were frequently noisy as a result of these adjustments and participant movement, and were therefore excluded when calculating variances and Hurst exponents, as well as when performing ICA and calculating artifact thresholds for localized component filtering.

